# Signal processing capacity of the cellular sensory machinery regulates the accuracy of chemotaxis under complex cues

**DOI:** 10.1101/2020.12.10.419572

**Authors:** Hye-ran Moon, Soutick Saha, Andrew Mugler, Bumsoo Han

## Abstract

Chemotaxis is ubiquitous in many biological processes, but it still remains elusive how cells sense and decipher multiple chemical cues. In this study, we postulate a hypothesis that the chemotactic performance of cells under complex cues is regulated by the signal processing capacity of the cellular sensory machinery. The underlying rationale is that cells *in vivo* should be able to sense and process multiple chemical cues, whose magnitude and compositions are entangled, to determine their migration direction. We experimentally show that the combination of TGF-β and EGF suppresses the chemotactic performance of cancer cells using independent receptors to sense the two cues. Based on this observation, we develop a biophysical framework suggesting that the antagonism is caused by the saturation of the signal processing capacity, but not by the mutual repression. Our framework suggests the significance of the signal processing capacity in the cellular sensory machinery.

## Introduction

Cell chemotaxis - the biased migration of cells toward a chemical cue - is a critical step in various pathological and physiological processes, including cancer metastasis, embryogenesis, and wound healing (Welch and Hurst, 2019). For multiple cell types, including cancer, cells are exposed to multiple signals within the complex chemical and physical environments during migration (Quail and Joyce, 2013; Roussos et al., 2011). These include multiple cytokines such as transforming growth factor-β (TGF-β), epidermal growth factor (EGF), stromal cell-derived factor-1α (SDF-1α), and CCL2 (Fernandis et al., 2004; Mantovani et al., 2002; Quail and Joyce, 2013; Roussos et al., 2011). Furthermore, the nature of the cytokine signals varies in space and time (Odenthal et al., 2016; Van Haastert and Devreotes, 2004). Cellular sensing is critical for deciphering the complex signals and accurately migrating in the direction of the physical and chemical cues whose temporal and spatial characteristics are tangled.

Prior studies on chemotaxis have focused on identifying molecules to establish the signal transduction network for a given chemical cue. Intracellular signaling networks involve a series of processes relevant to diverse cell physiological responses (Han and Lo, 2012; Ikushima and Miyazono, 2010). Research efforts to identify the associated signaling molecules and systematically construct their networks have advanced our understanding of how cells process the signals that result in a chemotactic response (Miyagawa et al., 2018; Odenthal et al., 2016; Roussos et al., 2011; Swaney et al., 2010). Quantitative modeling involving chemical kinetics or logical circuits has extensively contributed to constructing signaling networks in systems biology (Barberis and Verbruggen, 2017; Le Novère, 2015). These approaches elucidate the behavior of the cellular machinery, in which the physical and chemical components of the cell function together to carry out sensing, processing, and migration. At the same time, biophysical approaches have measured cellular sensing capacity in a quantitative manner (Hu et al., 2010; Thomas and Eckford, 2016; Varennes and Mugler, 2016). Physical limits to the precision of sensing have been investigated considering the biochemical nature of signal-receptor binding relations (Hu et al., 2010; Varennes and Mugler, 2016). The effect of receptor crosstalk in enhancing the precision of chemical sensing of multiple ligands has been demonstrated with quantitative modeling (Carballo-Pacheco et al., 2019). In cellular signal processing, intracellular protein diffusion and activation are critical to understanding the cellular sensory function (Soh et al., 2010). Not only the spatial variation, but also the temporal relations were used to develop a better understanding and physically predictable models (Gupta et al., 2020).

Cells are exposed to multiple cues whose chemical composition and physical nature are complex. Cellular capability to process the multiple signals has been addressed in functional crosstalk between either receptors or pathways (Hart et al., 2005; Lappano and Maggiolini, 2011; Shi and Chen, 2017; Wang et al., 2011). The EGFR cascade, an example of RTK, cross-communicates with the GPCR-mediated chemotaxis pathways in breast cancer (Hart et al., 2005). RTKs also interplay with transforming growth factor-β (TGF-β) through Smad-dependent transcription, or PI3K/AKT activation (Shi and Chen, 2017). The crosstalk at either the molecular or network level is reflected in cellular responses. Mosadegh et al. showed that EGF was cooperative in mediating breast cancer chemotaxis induced by SDF-1α gradient. (Mosadegh et al., 2008).In addition, Uttamsingh et al. (Uttamsingh et al., 2008) showed a synergistic effect of EGF and TGF-β in inducing invasion/migration capability in epithelial cells.

However, how cells transduce multiple signals is still poorly understood. This is a significant knowledge gap since cells *in vivo* need to process complex chemical cues to determine the migration direction. The question remains: Is there a limit to cellular capacity to process multiple signals, and if so, what happens when cells exceed it?

In this study, we investigate the effects of the signal processing capacity of the cellular sensory machinery on chemotaxis. Our central hypothesis is that chemotaxis performance is regulated by the signal processing capacity of the cellular sensory machinery. This hypothesis is tested by assessing chemotaxis of human breast cancer cells (MDA-MB-231) and murine pancreatic cells (eKIC) under single or combined cues of TGF-β1 (denoted as TGF-β) and EGF by use of a microfluidic platform. On the microfluidic platform, chemical cues of TGF-β and EGF are created with different gradient strengths and background concentrations. Then, a minimal signal processing machinery is modeled to address the experimental results. Using mathematical modeling, we hypothesize that the saturation of the pathways after they converge downstream limits the chemotactic performance. We further discuss the effects of signal processing on chemotaxis to lay the groundwork of a general theoretical model. The model based on the hypothesis reproduces the experimental results and predicts that the suppression of chemotactic performance can alternatively be caused by saturating the shared pathway with background signal strength. We confirm this prediction with further experiments on both cell lines. Our results constitute a biophysical framework of cellular response to multiple inputs and suggest that cellular capacity determines chemotactic accuracy under complex signals.

## Results

### The combination of TGF-β and EGF gradients is not synergistic for the chemotaxis of breast cancer cells

To understand the cell chemotactic response to multiple signals, we present gradient signals simultaneously using a microfluidic platform. The chemotaxis platform has been widely used to apply a chemical gradient to the cells embedded in the 3D extracellular matrix (Varennes et al., 2019; Wu et al., 2013). The platform is composed of three microfluidic channels, as shown in Figure. 1A. A center channel contains cells with a type I collagen matrix, supplementing nutrients from the two side channels of the source and sink filled with the basic culture medium. A chemical gradient is developed in the center channel by diffusion caused by a concentration difference between source and sink channels . We verify the concentration profiles by simulating the diffusion behaviors of TGF-β with 10kDa FITC-dextran (Figure. 1A). The concentration gradient is developed as a linear profile in the center channel, stably retained for more than 9 hours from 3 hours after perfusing FITC-dextran solution through the source channel while the sink channel is filled with FITC-dextran free medium. When we perfuse the FITC-dextran solution with the same concentration for both source and sink channels, the uniform concentration is quickly established in the platform (Figure. 1A).

**Figure. 1.**
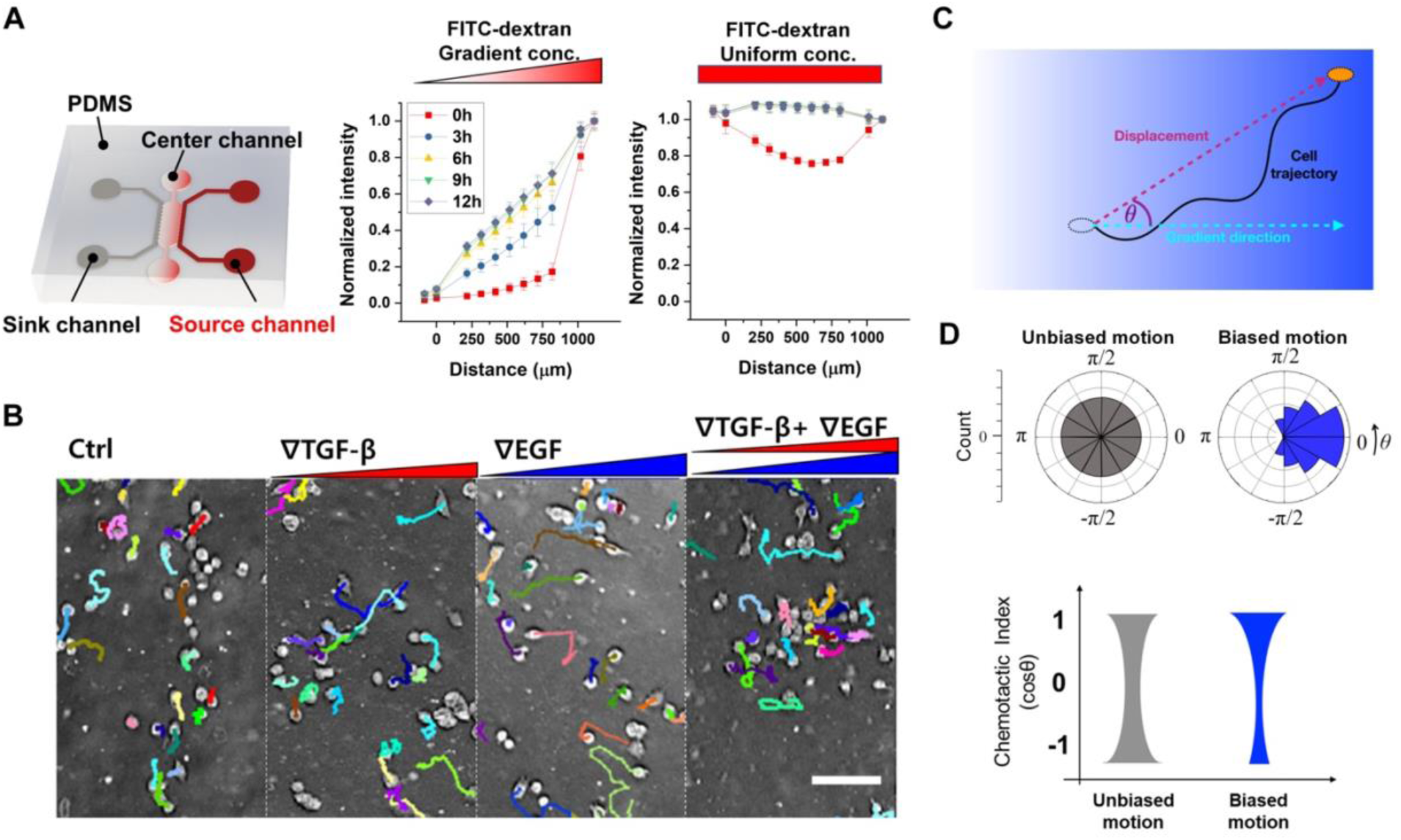
Chemotaxis platform and chemotaxis characteristics. (A) Schematic description of a microfluidic chemotaxis platform to induce the chemical gradient. 10kDa FITC dextran (simulating TGF-β) develops a linear profile when supplied at the sink channel, or a uniform profile when supplied at both channels in the chemotaxis platform. (B) Representative micrographs of MDA-MB-231 cells with their trajectories for 9 hours. (C) Characterization of the measured cell trajectory. (D) Schematics of angular (θ) distribution of cell trajectories and corresponding chemotactic index (CI) distributions of unbiased (gray) and biased motions (blue).

By using the chemotaxis platform, we expose the cells to single or combined cues, track cells’ migration, and analyze the chemotaxis characteristics of the cell trajectories to compute chemotactic index (CI) and speed (**Movie S1**, and Figure.S1). The CI is the cosine of the angle between a cell’s displacement and the gradient direction for a trajectory, indicating chemotactic accuracy toward the gradient (Figure. 1C). The CI can range between -1 and 1. Higher CI indicates more accurate chemotaxis in response to an attractant. CI = 1 means that the cell perfectly follows the gradient direction, whereas CI = 0 means that the cell is unbiased. The CI distribution of the unbiased group of cells is U-shaped with a median of 0. In contrast, cell trajectories showing biased motions exhibit a biased CI distribution toward 1 with a positive median (Figure. 1D).

The signal processing capacity of cells under multiple cues is anticipated to either synergistically or antagonistically affect the cell chemotactic performance. To elucidate the effect of combined cues on chemotaxis, we simultaneously expose both TGF-β and EGF gradients to the cells using the chemotaxis platform. We use a triple-negative breast cancer cell line, MDA-MB-231, recognized as a highly metastatic cell type (Chavez et al., 2010). Chemotactic response of MDA-MB-231 has been reported with several growth factors, including TGF-β and EGF (Varennes et al., 2019; Wang et al., 2004). Our previous study reported that MDA-MB-231 cells were significantly biased toward the TGF-β gradient direction (Varennes et al., 2019). Here, we investigate four groups: no growth factor (control), a 50nM/mm TGF-β gradient, an 800nM/mm EGF gradient, and the two gradients of TGF-β and EGF combined (Figure 1B; Figure. 2). In control, we see that the cell migration shows random movement with uniformly distributed migratory directions, whose CI distribution is U-shaped (Figure. 2C **gray**). On the other hand, the CI distribution for cells with either the TGF-β or EGF gradient shows a biased distribution toward 1 with significantly higher median CIs compared to the control (Figure. 2C **red and blue**). Surprisingly, the CI in response to the combined gradients decreases compared to the response to each individual gradient (Figure. 2C**, purple**), demonstrating that combining the two gradients has no synergistic effect on the CI.

**Figure. 2.**
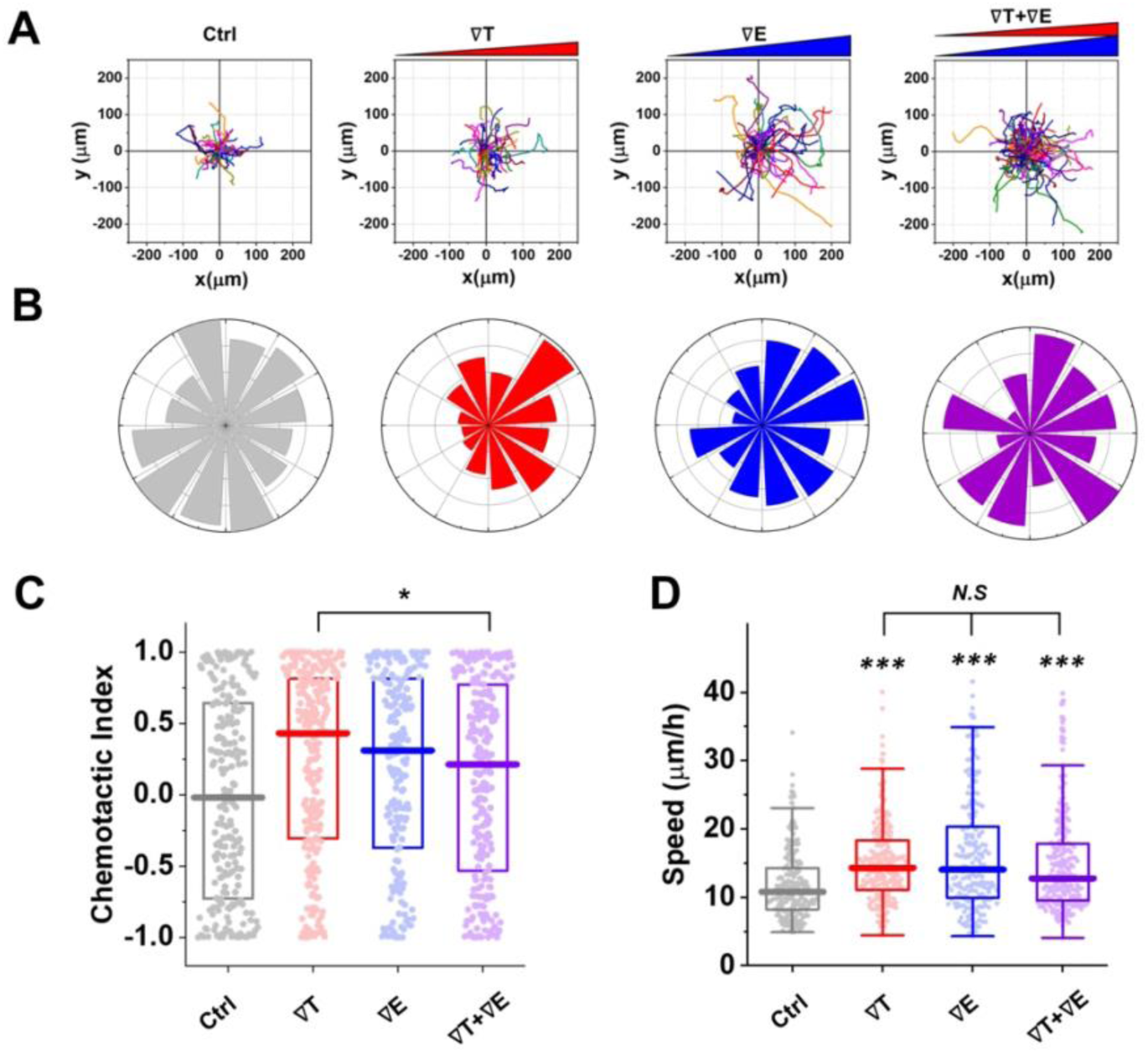
Chemotactic performance of breast cancer cells is not synergistically augmented when TGF-β and EGF gradients are simultaneously imposed. (A) Cell trajectories of a representative sample for control, 50nM/mm TGF-β gradient (∇T), 800nM/mm EGF gradient (∇E), and combined gradients of 50nM/mm TGF-β with 800nM/mm EGF (∇T+∇E) collected for 9 hours. Trajectories in a sample include >35 cells. (B) Angular (θ) distribution of cell trajectories from a representative sample for control (gray), ∇T(red), ∇E(blue), and ∇T+∇E(purple). (C) Distribution of chemotactic index (CI) from all trajectories collected from experimental trials (N>3) including 158-233 trajectories per condition. Box represents quartiles with a median line in the middle of the box. (D) Distribution of speed with a median line from all collected trajectories. Box: interquartile range (IQR) ± 1.5 IQR whiskers. A dot represents data from a single trajectory. (***: p<0.001, N.S: no significance with p>0.05 in Mann-Whitney test) (See also Figure S1.)

It is a remarkable result since many studies have reported the functional cooperation of multiple signals rather than the antagonism that we see here (Pang et al., 2016; Uttamsingh et al., 2008). In fact, cells lose their chemotactic ability if the gradient strength is not sufficient. However, we observe similar CI values when either TGF-β or EGF gradient is applied individually, even though the gradient strengths are different from each other. This suggests that they are appropriately tuned to their corresponding receptors. The observed lack of synergy is surprising also because TGF-β and EGF bind to independent receptors on the cell surface: TGF-β binds to the TGF-β receptors, which are transmembrane serine/threonine kinase complexes composed of type 1 and type 2 subunits (Ikushima and Miyazono, 2010); whereas EGF binds EGFR which is a receptor tyrosine kinase superfamily (Han and Lo, 2012). Thus, the combined effect of TGF-β and EGF reflects cellular signal processing, excluding an interplay in a receptor-binding level.

Interestingly, cells show significantly enhanced speed compared to control in both the individual gradient cases and under the combined gradients (Figure. 2D). In fact, the cell speed is not significantly enhanced with the combined gradients compared to the individual gradient cases. This demonstrates that combining the gradients of two chemoattractants does not have a synergistic nor antagonistic effect on the cell speed.

### The multitasking response is not cell-type specific

To evaluate the cell-type dependency in cellular multitasking, we investigate multiple-cue chemotaxis in pancreatic cancer cells. eKIC is used in this study, which is a murine pancreatic cancer cell line. eKIC is driven by the genetically engineered mouse model having Kras and p16 mutations showing epithelial phenotype (Bradney et al., 2020; Seeley et al., 2009). The cell line is known to be responsive to TGF-β, showing invasion features (Bradney et al., 2020). First, we evaluate the migratory behaviors of eKIC in the chemotaxis platform as a control, excluding any growth factors. The trajectories show an average CI close to 0, as shown in Figure. 3 (gray). Then, we expose a single cue of either a 10 nM/mm TGF-β gradient or a 200 nM/mm EGF gradient. We find that the eKICs are highly responsive to both TGF-β gradient and EGF gradient (Figure. 3, red and blue). Indeed, the median CI values are significantly enhanced in both gradient groups compared to the control group. Interestingly, we observe the antagonism in eKIC in the presence of the combined TGF-β and EGF gradients: the CI is significantly lower than the CIs of both groups of a single gradient (Figure. 3C, purple). The speed is enhanced significantly compared to the control (Figure. 3D), and again the speed under combined TGF-β and EGF gradients does not show significant differences to the speeds in single gradient groups, neither TGF-β nor EGF gradient. These results indicate that the lack of synergy in the chemotactic performance is not cell-type-specific responses. Instead, the present results suggest that the cell’s multitasking response might be governed by physical principles rather than cell-type-specific details.

**Figure. 3.**
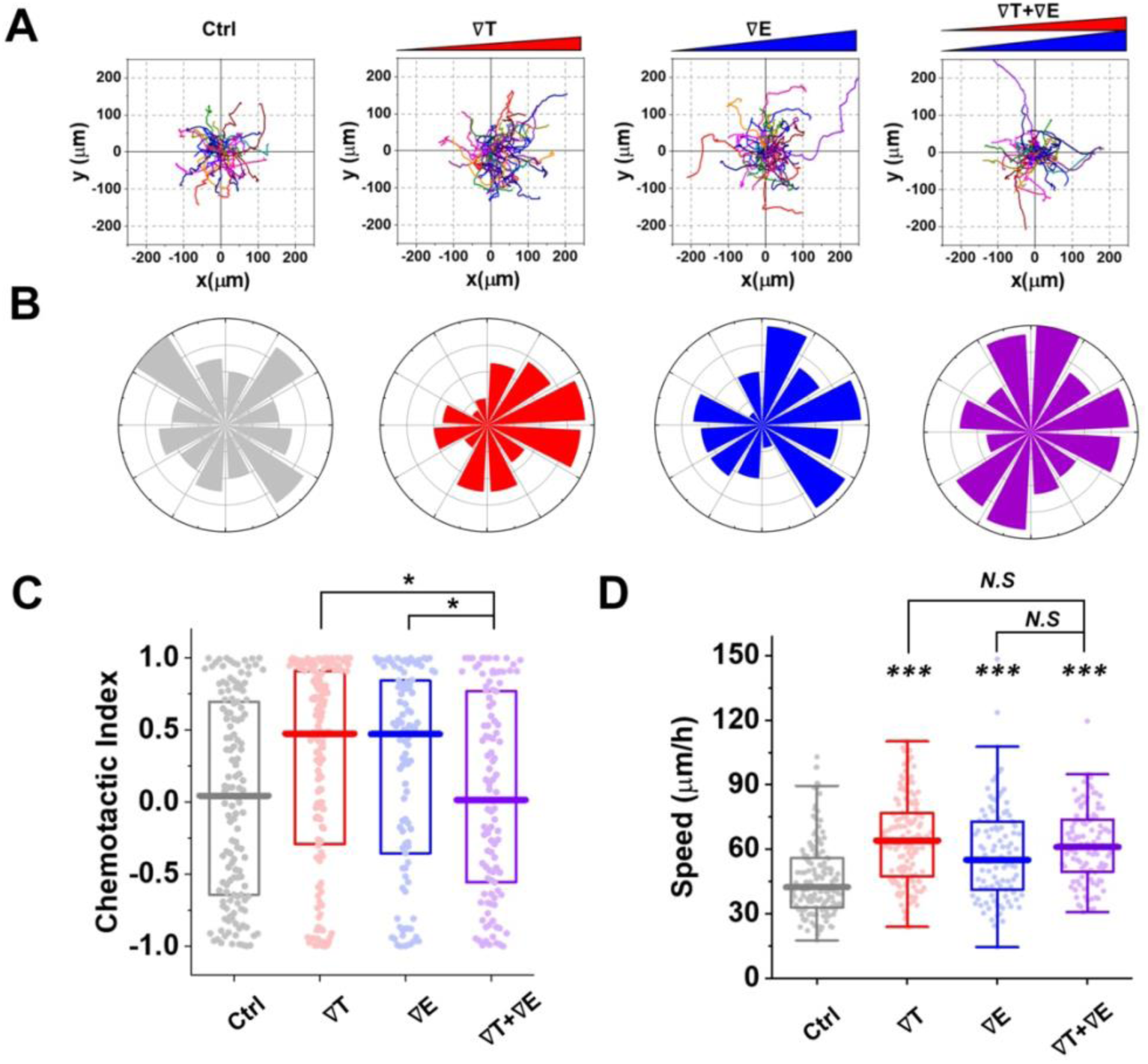
Antagonism of pancreatic cancer cells in the combined signal environment. (A) Cell trajectories of a representative sample for control, 10nM/mm TGF-β gradient (∇T), 200nM/mm EGF gradient (∇E), and combined gradients of 10nM/mm TGF-β with 200nM/mm EGF (∇T+∇E) collected for 3 hours. Trajectories in a sample include >35 cells. (B) Angular (θ) distribution of cell trajectories from a representative sample for control (gray), ∇T(red), ∇E(blue), and ∇T+∇E(purple). (C) Distribution of chemotactic index (CI) from all trajectories collected from experimental trials (N>3) including >100 trajectories per condition, respectively. Box represents quartiles with a median line in the middle of the box. (D) Distribution of speed from all collected trajectories. Box: interquartile range (IQR) ± 1.5 IQR whiskers with a median line. A dot represents data from a single trajectory. (***: p<0.001, N.S: no significance with p>0.05 in Mann-Whitney test) (See also Figure S1.)

### The multitasking response cannot be explained by mutual repression but can be explained by a shared pathway

Because TGF-β and EGF bind to different receptors (Han and Lo, 2012; Ikushima and Miyazono, 2010), the fact that combining both signals suppresses the CI is not due to receptor saturation, but rather due to the signaling network downstream. To understand this effect, we turn to mathematical modeling. We suppose that the migratory response is ultimately governed by an internal molecular species *M*. In order for the TGF-β or EGF gradient to be translated into biased migration, there must be a difference in the concentration of *M* between the front and back halves of the cell (Ellison et al., 2016). Therefore, we take this difference Δ*m* to be a proxy for the CI (Figure. 4A). We have seen that the cell speed increases significantly in the presence of either TGF-β or EGF (Figure. 2D and 3D). Therefore, we take the average concentration of *M* in both halves of the cell, *m*, to be a proxy for speed (Figure. 4A). In principle, the speed could also depend on the bias Δ*m*. However, we have checked experimentally that cell speed for a given background level of TGF-β or EGF is not significantly different whether the corresponding gradient is present or not (Figure. S2). Thus, we take speed to be a function of *m* alone. It now remains to propose a model of the signaling network connecting TGF-β and EGF to *M* that can explain the data.

**Figure. 4.**
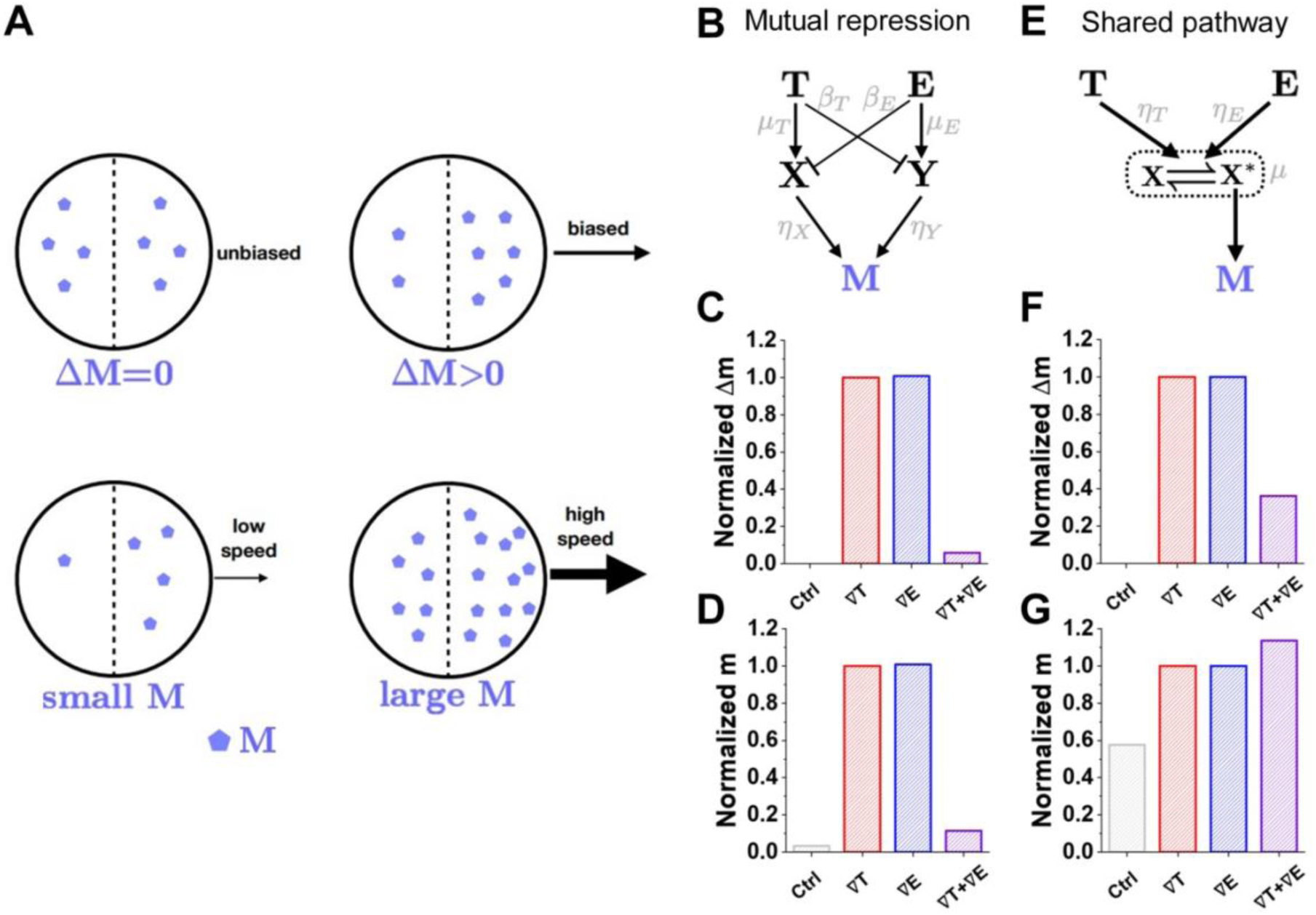
Mathematical model explains antagonism by saturation of a shared pathway. (A) In the model the cell speed and chemotactic index (CI) scale with the concentration *m* of an intracellular species *M*, and its concentration difference Δ*m* between the front and back of the cell, respectively. (B) In the mutual repression model, TGF-β and EGF mutually repress the other’s activation pathway. (C) Δ*m* exhibits antagonism (the response to both gradients is smaller than that of either alone). (D) *m* also exhibits antagonism. (E) In the shared pathway model, TGF-β and EGF convert a common component to its active state. (F) Δ*m* exhibits antagonism. (G) *m* does not exhibit antagonism. Only the shared pathway results are consistent with the data in Figures. 2 and 3. C, D, F, and G each shows results averaged over parameter space (see **Method details**) and normalized by the TGF-β gradient case (red). (See also Figure S2.)

A straightforward hypothesis for the observed antagonism in the CI is the presence of mutual repression: TGF-β activates *M* through pathway one, and EGF activates *M* through pathway two, but TGF-β also represses pathway two and vice versa. A minimal example of such a mutually repressive network is shown in Figure. 4B. Here we have introduced species *X* and *Y*, which are the targets of the repression, and dimensionless parameters *η_X_* ,*η_Y_* , *μ_T_* , *μ_E_* , *β_T_* , and *β_E_* which define the strength of the edges. Writing down the rate equations for this network and solving for the steady-state, we find

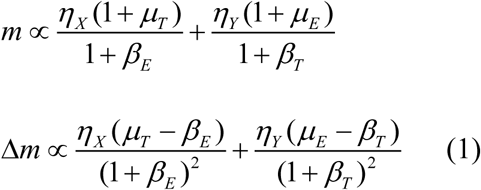

where the dimensionless parameters are defined in terms of the reaction rates (see **Method details**). These expressions apply to the case where both signals are present, while specific limits apply to the case where only the TGF-β gradient is present ( *μ_E_* = *β_E_* = 0 ), the case where only the EGF gradient is present ( *μ_T_* = *β_T_* = 0 ), and the control ( *μ_T_* = *β_T_* = *μ_E_* = *β_E_* = 0 ).

To investigate whether this model can explain the data in Figures. 2 and 3, we focus on parameters for which Δ*m* is smaller when both gradients are present than when either is present alone (Figures. 2C and 3C). The result, averaged over all such parameters (see **Method details**), is shown in Figure. 4C, and we see that this model can explain an antagonistic effect on the CI. However, for these same parameters, we also see an antagonistic effect on *m* in Figure. 4D, which is inconsistent with the speed data in Figures. 2D and 3D. The reason that antagonism for Δ*m* and *m* coincide in this model is that the mutual repression suppresses the front-back bias in the output molecule *M* by suppressing the production of *M* itself. We conclude that crosstalk cannot explain the data.

We, therefore, put forward an alternative hypothesis that the antagonism in the CI is due to a shared signaling pathway. Specifically, if the pathways by which TGF-β and EGF activate *M* converge at a certain point, it could be that the shared pathway is saturated when both signals are present, but not when each signal is present alone. A minimal example of such a network is shown in Figure. 4E. Here *X* is the shared component, and TGF-β and EGF catalyze the reversible conversion of *X* to an activated state *X*\*, which produces *M*. The presence of a convertible species introduces the possibility of saturation, where all *X* molecules have been converted to the *X*\* state, and no further conversion can occur. The dimensionless parameters *η_T_* and *η_E_* define the strengths of the catalysis reactions, and *μ* defines the degree to which the conversion reaction is unsaturated (*μ* → 0 ) or saturated ( *μ* → 1). Again, writing the rate equations and solving for the steady-state, we find

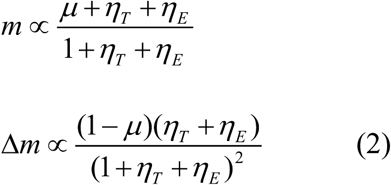

where *η_T_* ,*η_E_* and *μ* are defined in terms of the reaction rates, and the limiting cases are the same as those in Eq. 1. Again, averaging over all parameters for which Δ*m* is smaller with both gradients than with either alone (Method details), we see that this model can explain both antagonism for the CI (Figure. 4F) and a lack of antagonism for the speed (Figure. 4G), consistent with the data in Figures. 2 and 3. The reason that antagonism occurs for Δ*m* but not *m* in this model is due to the saturation: as all *X* molecules become converted to the *X*\* state, the difference in the concentration of *M* molecules from front to back vanishes, while the total concentration of *M* molecules remains high. The saturation causes loss of synergy in the value (*m*), but it causes antagonism in the slope (Δ*m*). Because gradient sensing depends on the slope (Δ*m)*, a value-saturating function causes slope-antagonism, which indicates a saturation of signal processing capacity. This can be understood by thinking about what happens to Δ*m* when *m* is close to saturation. In our system when only one gradient is present, *m* is away from saturation, so there is a large Δ*m*. But if both gradients are applied simultaneously, then *m* is close to the saturating value, and Δ*m* is small. Because the response depends on the difference Δ*m* in molecule number between the front and back of the cell, the decrease in Δ*m* is sufficient to explain the antagonism behavior in cells. We conclude that the saturation of a shared pathway can explain the data.

### The shared pathway model successfully predicts the cellular response to elevated signal background

The key mechanism behind the shared pathway model is that the presence of a second gradient signal brings the shared pathway too close to its saturation point and therefore reduces the chemotactic bias while not reducing the speed. Given this mechanism, the model makes an important prediction: the second signal need not be graded to bring the shared pathway to its saturation point. Indeed, we see in Figure. 5 that if the TGF-β gradient is combined with a uniform EGF background alone, then the bias Δ*m* is reduced (Figure. 5A, black), whereas the average *m* is not (Figure. 5B, black). In fact, the model also makes a second prediction: the elevated background need not even come from the other signal; it could come from the same signal. Indeed, we also see that if the TGF-β gradient is combined with an additional uniform TGF background, then the bias Δ*m* is reduced (Figure. 5A, maroon) whereas the average *m* is not (Figure. 5B, maroon).

**Figure. 5.**
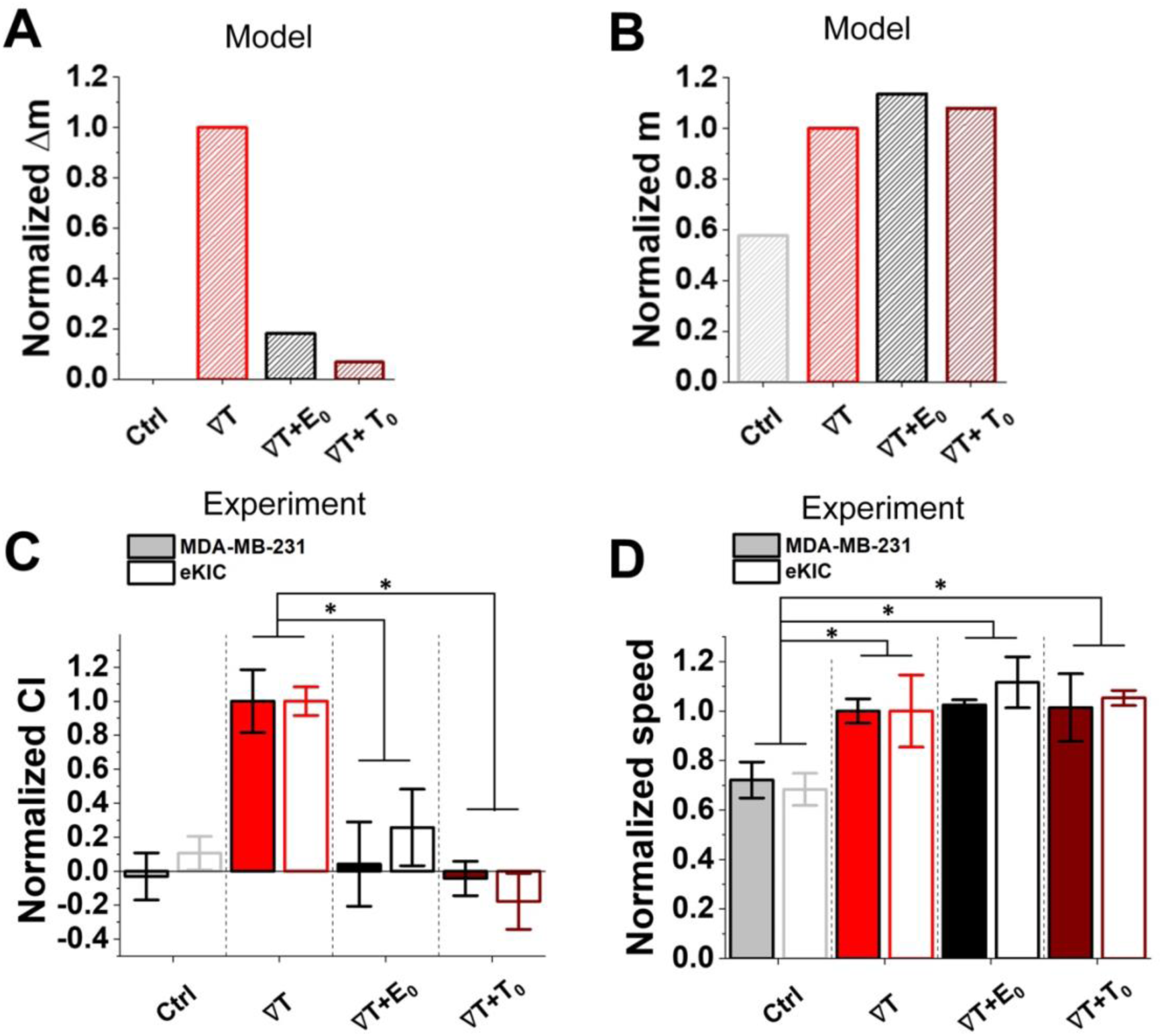
Model predicts and experiments confirm that elevated signal background reduces chemotactic bias but not speed. The model predicts that (A) the bias Δ*m* decreases when a TGF-β gradient is combined with a uniform background of either EGF or TGF-β, whereas (B) the average *m* does not. (See also Figure S3.) Experiments confirm that for either cell type, (C) the chemotactic index (CI) is significantly suppressed when the TGF-β gradient is combined with a uniform background of either EGF or TGF-β, whereas (D) the speed is not, consistent with the predictions. (*: p<.05, **: p<.01 in Student t-test). In all panels, the values are normalized by the TGF-β gradient case (red). The model uses Eq. 4 with Δ*e* = 0 (EGF background) or {*e* = 0, Δ*e* = 0, *t* → 10*t*} (TGF background) and the same parameters as in Figure. 4. The experiment uses the same TGF-β gradient as in Figures. 2 and 3 (50 nM/mm for MDA-MB-231, 10 nM/mm for eKIC) combined with either uniform EGF (400 nM for MDA-MB-231, 100 nM for eKIC) or uniform TGF-β (200 nM for MDA-MB-231, 50 nM for eKIC). In experiment, bar represents mean of the medians ± S.E (n≥3). The medians are collected from >35 trajectories in a sample, respectively. (*: p<0.05, **: p<0.01, and N.S: no significance with p>0.05 in student’s t-test) (See also Figure S1.)

To test these predictions, we expose each cell type to its respective TGF-β gradient as in Figures. 2 **and** 3, combined with a uniform EGF background (equal to the EGF concentration in the center of the device in Figures. 2 **and** 3). We see that in both cases, the CI is significantly suppressed (Figure. 5C, black), whereas the speed is not (Figure. 5D, black), as predicted. Then, we expose each cell type to the TGF-β gradient combined with a uniform TGF-β background concentration. Again, we see that in both cases, the CI is significantly suppressed (Figure. 5C, maroon), whereas the speed is not (Figure. 5D, maroon), as predicted. These results support the hypothesis that the growth factors are saturating a shared signaling pathway, and therefore that the capacity of the cell to multitask is being exceeded.

Notably, the mutual repression model makes predictions in these two cases that are distinct from those of the shared pathway model and, therefore, inconsistent with the data (Figure. S3).

## Discussion

We have investigated cellular multitasking capacity in the context of the migratory response to simultaneous growth factor gradients in a 3D microfluidic environment. Surprisingly, for two different cell types, we have found that the chemotactic bias is suppressed below that for either growth factor alone, whereas the cell speed remains high. Because the growth factors bind to separate receptors, the suppression is not a consequence of simple receptor saturation, rather must result from the signaling network downstream. Using mathematical modeling, we have found that the suppression cannot be explained by a network with mutually repressive crosstalk but can be explained by the convergence to a shared pathway. Our shared pathway model has predicted that the chemotactic bias, but not speed, can be alternatively suppressed by elevating the background level of either growth factor, which our experiments have confirmed. Our results emphasize the fact that the quantitative features, not just the topological features of a signaling network, are necessary to understand a behavioral response, especially a counterintuitive response such as antagonism.

The model successfully predicts the cellular response regardless of cell type. This suggests that chemotaxis is critically moderated by not only specific environmental cues and cell-type-specific biochemistry, but also the intrinsic cell capability to process information within an inherently limited channel. Indeed, genetic mutations commonly observed in cancer cells strongly affect the migration by promoting the invasion process via specific signaling pathways such as Snail and Twist (Muller et al., 2011). Yet, the present results show that the cell response could be predicted from our framework regardless of the cancer type, implying that cellular multitasking capacity is not a cancer type-specific signaling effect.

The antagonistic response to two signals in the model results from the saturation of a shared pathway. In this saturating regime, we have seen that the model explains the experimental data and makes predictions that our further experiments confirmed. Yet, the model also predicts more conventional synergistic behavior outside the saturating regime. Specifically, defining an amplification factor *α* = *η_T_* +*η_E_* and a relative pathway strength *ρ* = *η_T_*/*η_E_* in terms of the parameters of Eq. 2, we see in Figure. 6A that the model supports both antagonistic and synergistic regimes (see **Method details**). Intuitively, antagonism arises when each individual pathway is amplified ( *α* >>1) and the two pathways are balanced ( *ρ □* 1 ), as illustrated in Figure. 6B. In contrast, synergy arises when the amplification is not as strong, as then the shared pathway is not saturated, and the capacity of the channel is not exceeded.

**Figure. 6.**
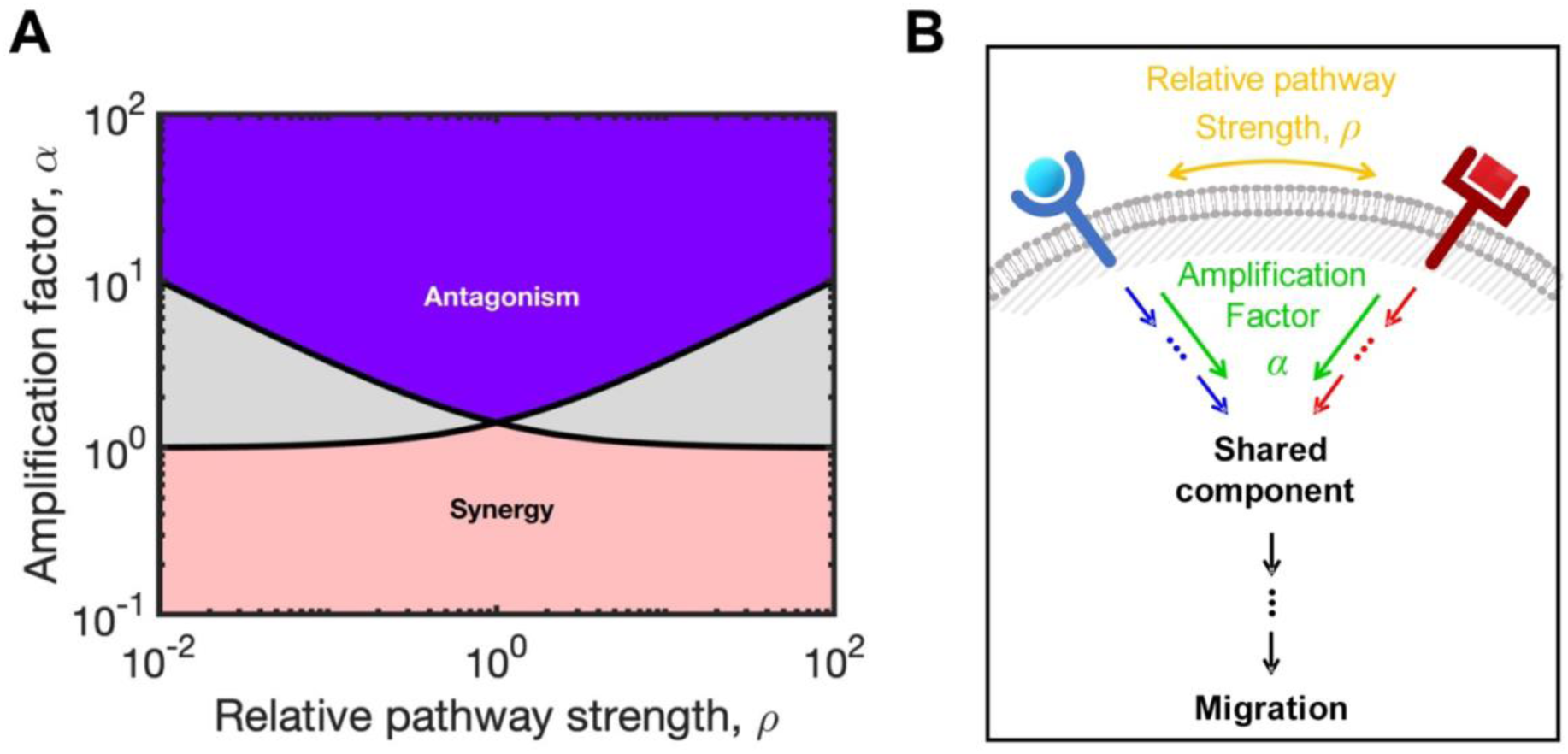
Schematic of cellular multitasking capacity for chemotaxis. (A) The shared pathway model predicts antagonism in the chemotactic response (as in the experiments) for large amplification factor *α* and balanced relative pathway strength *ρ*, but synergy in the response for small *α* (see **Method details** for phase boundaries). (B) Illustration of *α* and *ρ* in the general context of a signaling network defined by pathway convergence.

The fact that the proposed signaling mechanism supports either antagonism or synergy may help explain why previous work has observed synergistic effects between growth factors, including TGF-β and EGF. Both of these growth factors have been identified to regulate cell invasiveness and motility (Hou et al., 2012; Moustakas and Heldin, 2008; Roussos et al., 2011), with TGF-β in particular known to promote epithelial-mesenchymal transition, which is a critical step in cancer invasion (Ikushima and Miyazono, 2010; Moustakas and Heldin, 2008). In fact, TGF-β is generally considered a cooperative effector when it combines with other growth factors (Pang et al., 2016; Uttamsingh et al., 2008). Most notably, Uttamsingh, et al. reported that the roles of TGF-β- and EGF-mediated signaling pathways combine synergistically in inducing cell invasion and migration (Uttamsingh et al., 2008). However, it is important to note that this study focused on motility and invasiveness, not directional bias, and indeed here, we do not observe an antagonistic effect on cell speed. This comparison emphasizes the need to distinguish directional and non-directional measures of cell motility, as well as the difference between cell-intrinsic motility factors and the physical aspects of the signaling environment (Clark and Vignjevic, 2015; Endres and Wingreen, 2008; Varennes et al., 2019). Our discovery of antagonism by the cellular capacity suggests the necessity of a quantitative approach to multiple-cue cell migration studies.

Both TGF-β and EGF are secreted by stromal cells, which typically surround the solid tumors. (Bulle and Lim, 2020; Roussos et al., 2011; Shibue and Weinberg, 2017) Thus, we anticipate that these growth factors are usually imposed along with the aligned directions. However, considering the spatial and temporal heterogeneity of tumor tissue structure, it may depend on the tumor structure and development stage. The other scenarios for the gradient alignments could be further investigated with the controllable microfluidic platform, for example, by imposing conflicting gradients. Nonetheless, the present study provides insight into how combined gradients of multiple growth factors in the tumor microenvironment could affect cancer cell migration behavior.

Our framework proposes that two receptor-activated signaling pathways converge to a single pathway to promote migration (Figure. 6B), and there is ample biochemical evidence that this structure is plausible. From a general standpoint, although many signaling pathways regulate chemotaxis (Charest and Firtel, 2007; Chung et al., 2001; Gandalovičová et al., 2016; Stuelten et al., 2018), all must converge to activate a set of common cellular responses, including actin polymerization and organization, cytoskeleton dynamics, and adhesion (Roussos et al., 2011; Swaney et al., 2010; Van Haastert and Devreotes, 2004). More specifically, complex signaling pathways are known to converge onto central functional molecules (idealized in our model by the species *X*) such as PIP_3_ in *Dictyostelium*, cofilin in breast carcinoma cells (Swaney et al., 2010), Rho GTPases, Smad-dependent transcription, or PI3K/AKT activation cascades (Charest and Firtel, 2007; Chung et al., 2001; Corallino et al., 2018; Hart et al., 2005; Lappano and Maggiolini, 2011; Li et al., 2013; Miyagawa et al., 2018; Shi and Chen, 2017; Swaney et al., 2010).

Our results suggest that the antagonistic response to two gradient signals in these cells is due to saturation of the sensory pathway. However, well-characterized networks in other sensory systems, such as the bacterial chemotaxis network, avoid saturation by adapting their operating point to the background concentration (Alon et al., 1999; Mello and Tu, 2007). This adaptation relies on a timescale separation between effector phosphorylation and receptor methylation that, for whatever reason, may be difficult to achieve in the cells studied here. Alternatively, the growth factor concentrations at which saturation sets on in our system may be outside of the physiological range to which these cells are typically exposed. Further characterization of the sensory network is needed to shed light on the question of adaptation in these cells.

## Limitations of study

Although the biophysical framework is capable of illustrating the cellular multitasking capacity shown in the experiments, further research is warranted. The present framework may limit the explanation regarding a mode of migration. The physical constraints driven by the extracellular matrix (ECM) regulates features of cell migration as well. Since the chemotactic response and the relevant signaling pathways inducing chemotaxis could be varied depending on the mode of the migration (Clark and Vignjevic, 2015; Gandalovičová et al., 2016), the model may be improved further by considering migration modes highly regulated by physical confinement.

## Acknowledgments

This work was partially supported by grants from the National Institutes of Health (R01 CA 254110 and P30 CA023168). Publication of this article was funded in part by Purdue University Libraries Open Access Publishing Fund.

## Author contributions

AM and BH conceived the idea. HM and SS performed the research and acquired the data. All authors discussed the results and wrote the manuscript.

## Inclusion and diversity

We worked to ensure diversity in experimental samples through the selection of the cell lines. The author list of this paper includes contributors from the location where the research was conducted who participated in the data collection, design, analysis, and/or interpretation of the work.

## STAR METHODS

### KEY RESOURCE TABLE

Chemicals and recombinant proteins

Experimental models: Cell lines

Software and algorithms

### RESOURCE AVAILABILITY

#### Lead contact

Further information and requests for resources and reagents should be directed to and will be fulfilled by the lead contact, Bumsoo Han.

#### Materials availability

This study did not generate new unique reagents.

#### Data and code availability

All data reported in this paper will be shared by the lead contact upon request.

All original code has been deposited at Zenodo and is publicly available as of the date of publication. DOIs are listed in the key resources table.

Any additional information required to reanalyze the data reported in this paper is available from the lead contact upon request.

## EXPERIMENTAL MODEL AND SUBJECT DETAILS

### Cell lines

A human breast cancer cell line (MDA-MB-231) and a murine pancreatic cell line (eKIC) were used in this study. MDA-MB-231 cells (ATCC, VA, USA) were maintained in Dulbecco’s Modified Eagle Medium/Ham’s F-12 (Advanced DMEM/F-12, Lifetechnologies, CA, USA) supplemented with 5% v/v fetal bovine serum (FBS), 2 mM L-glutamine (L-glu), and 100 µg ml^−1^ penicillin/streptomycin (P/S). eKIC was obtained from Dr. Murray Korc’s laboratory at the Indiana University. eKIC was isolated from KIC mice, which were established as a genetically engineered mouse model of PDAC (GEM). For the KIC mice, *Kras* was combined with the deletion of the *Ink4a* locus (*Ink4a/Arf^L/L^*) to generate the *Pdx1-Cre;LSL-Kras^G12D^;Ink4a/Arf^-/-^* GEM (Sempere et al., 2011; Whipple et al., 2011). The eKIC cells were cultured in RPMI 1640 with 2.05mM L-glutamine (GE Healthcare Bio-Sciences Corp., MA, USA) supplemented by 5% v/v fetal bovine serum (FBS) and 100 µg ml^−1^ penicillin/streptomycin (P/S). Both MDA-MB-231 and eKIC cells were regularly harvested by 0.05% trypsin and 0.53mM EDTA (Lifetechnologies, CA, USA) when grown to ∼80% confluency in 75 cm^2^ T-flasks and incubated at 37°C with 5% CO_2_. The harvested cell suspensions were used for experiments or sub-cultured. Both cells were maintained below 15^th^ passage while regularly kept in cryopreservation.

## METHOD DETAILS

### Chemotaxis assay

For the chemotaxis assay, cells were implanted in the chemotaxis platform, which is an *in vitro* microfluidic device developed to engineer the chemical environment.as shown in Figure. 1A. The chemotaxis platform was designed for engineering the chemical environment surrounding the cells embedded in the 3D extracellular matrix. The platform is composed of three microfluidic channels with 100μm in thickness. A center channel of 1mm wide, which aims to contain cells with a collagen matrix, is located in between source (top) and sink (bottom) channels with 300μm wide. At the end ports, the source and sink channels are connected to large reservoirs so that the culture medium can be supplemented through the channels. For the cell culture, the basic culture medium is filled in both source and sink channels. In order to develop a concentration gradient in the center channel, growth factor solution (TGF-β or EGF) based on the medium is added through the source channel. On the other hand, the sink channel is filled with growth factor-free medium. Assuming that there is neither flow nor any pressure differences between the channels, the concentration of the given growth factor (*i*) can be illustrated by the conservation equation of chemical species:

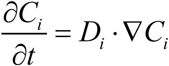

At the steady state, the concentration profile in the center channel goes to a linear. The linear profile can persist for a while with an assumption that is the concentration at the source and side channels are constant with a large volume of reservoirs. Consequently, cells cultured in the collagen matrix are exposed to a linear gradient of a specific soluble factor. The detailed technique to develop the microfluidic device and the diffusion principle in the platform was described in our previous study (Varennes et al., 2019).

Cells were embedded uniformly to type I collagen mixture (Corning Inc., NY, USA) supplemented with 10X PBS, NaOH, HEPE solution, FBS, Glu, P/S, and cell-culture level distilled water. Initial cell density was consistently 5×10^5^ cells/ml for MDA-MB-231 and 8×10^5^ cells/ml for eKIC respectively in the 2mg/ml type I collagen mixture. The cell-collagen mixture was loaded in the center channel of the microfluidic device. After loading, the cells in the collagen matrix were cultured with basic mediums for 24 hours. MDA-MB-231 cells were then exposed by serum-reduced medium for another 24 hours. The serum-reduced medium was supplemented by 1% v/v FBS instead of 5% v/v FBS in the basic medium. Due to a critical viability change in the serum-reduced culture condition (data is not shown), serum starvation was not conducted for eKIC. Then, cells were exposed by concentration gradients of growth factors, either Transforming growth factor beta-1 (TGF-β1, Invitrogen, CA, USA) or epidermal growth factor (EGF, Invitrogen, CA, USA).

### Characterization of cell migration

Live-cell imaging technique with time-lapse microscopy was used to characterize cell migration. An inverted microscope (Olympus IX71, Japan) was equipped with a stage top incubator as described in (Varennes et al., 2019), which maintain the microfluidic chemotaxis platform at 37°C with 5% CO_2_ environment during imaging. MDA-MB-231 cells in the chemotaxis platform were captured every 15 minutes for 9 hours. eKIC cells were captured every 5 minutes for 3 hours. The temporal intervals and durations for the time-lapse imaging were optimized for each cell line by considering the cell motility, ∼12μm/h for MDA-MB-231 and ∼50μm/h for eKIC in control, respectively (Figure. 2D **and** 3D). In both cases, the time-lapse imaging was started 3 hours after applying the growth factors for sufficient adjusting time to develop gradient profiles accordingly (Figure. 1B). The bright-field images were further processed to analyze cell migration. The cell area was defined by using the contrast differences between cells and background and converted to monochrome images by using ImageJ. A cell trajectory was illustrated as a collection of the centroid positions of cell areas at different time points. In tracking the cell movement, we excluded the cells undergoing division to avoid extra effect for cell polarity (Harley et al., 2008). Also, we excluded the stationary cells defined when the cells moved less than their diameter.

Biased cell motion is commonly characterized by directional accuracy, persistence, and motility (Endres and Wingreen, 2008; Roussos et al., 2011; Skoge et al., 2014; Varennes et al., 2019). In our previous study (Varennes et al., 2019), we have shown that the directional accuracy is a dominant metric in the chemotaxis of the 3D cultured cells migrating in ECM whereas cell persistence is barely changed by the chemical gradient. In this study, we therefore focus on accuracy and motility, not persistence.

We measure directional accuracy y with the commonly used chemotactic index (CI) (Endres and Wingreen, 2008; Skoge et al., 2014)

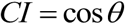

where *θ* is the angle between the net displacement of a trajectory and the gradient direction. The displacement is defined as a straight line connecting the initial and final points of a trajectory where we measure each trajectory for 9 hours for MDA-MB-231 and 3 hours for eKIC (Figure. S1B). In an experimental trial, multiple cells’ CI values are distributed throughout the range of -1 to 1 due to variations of cell response to the attractant. The CI distribution is U-shaped without any attractant, indicating that the cell migration directions are uniformly distributed; in contrast, the CI distribution with an attractant shows a biased distribution toward 1 (Figure. 1D) (Varennes et al., 2019).

Cell motility is quantified as an instantaneous speed along the cell trajectory:

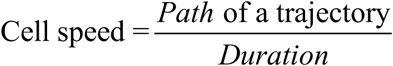

where the cell path is taken from a trajectory where measurement is taken every Δ*t* = 15 minutes.

### Defining the mathematical models and deriving the equations in the main text

For the mutual repression network in Figure. 4B, denoting the concentrations of TGF-β, EGF, *X*, and *Y* as *t*, *e*, *x*, and *y*, respectively, the rate equations are

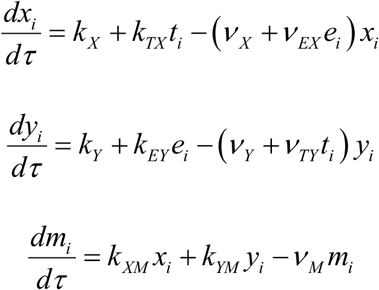

where *i* = 1, 2 denotes the front or back half of the cell, *τ* is time, *k* and *ν* denote production and degradation rates, respectively, and activation and repression are modeled as a linear dependence of production and degradation on the regulator, respectively. In steady state, the solution takes the form *m_i_* = *f* (*t_i_* ,*e_i_* ), where *f* follows algebraically from the rate equations. Because the changes in TGF-β and EGF concentrations across the cell, Δ*t* = *t*_2_ − *t*_1_ = *R*∇*T* and Δ*e* = *e*_2_ − *e*_1_ = *R*∇*E* , are much smaller than the average concentrations, *t* = (*t*_1_ + *t*_2_ )/2 and *e* = (*e*_1_ + *e*_2_ )/2 , for all but the cells that are within a few cell radii *R* from the sink channel of the device, the average *m* = (*m*_1_ + *m*_2_ )/2 and difference Δ*m* = *m*_2_ − *m*_1_ can be approximated as

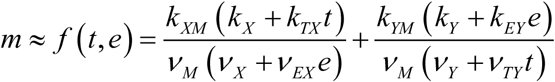

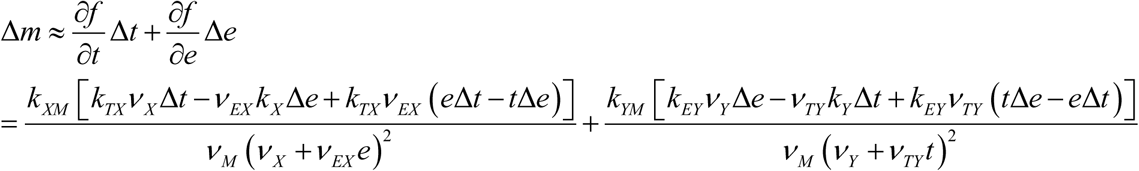

We nondimensionalize these expressions by defining the concentration scale

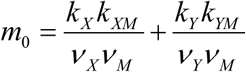

and dimensionless ratios

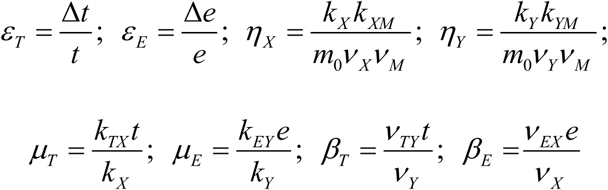

with which they become

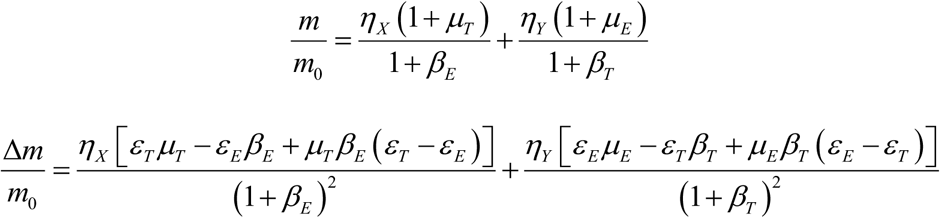

Because both the TGF-β and EGF concentrations are zero at the sink channel (without the elevated backgrounds), we have *t* = *d*∇*T* , Δ*t* = *R*∇*T* , *e* = *d*∇*E* , and Δ*e* = *R*∇*E* , where *d* is the distance of the cell from the sink, and *R* is the cell radius. Therefore, we see that *ε_T_* =*ε _E_* ≡ *ε* , and the previous equation simplifies to

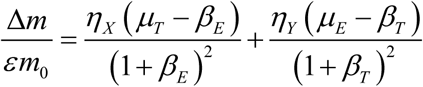

These expressions for *m* and Δ*m* are reproduced in Eq. 1 of the main text.

For the shared pathway network in Figure. 4E, the rate equations are

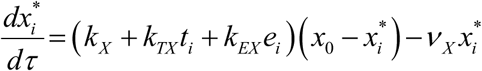

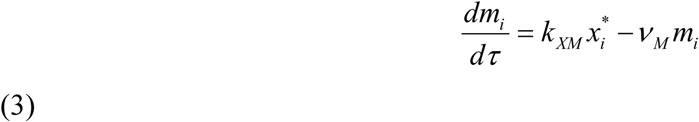

where *x*_0_ is the total concentration of *X* and *X*\* molecules, which we assume to be constant and equal in both halves of the cell. In steady state we have

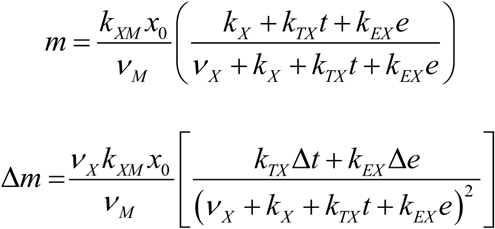

where we have applied the same approximations as in the mutual repression model. We nondimensionalize these expressions by defining the concentration scale

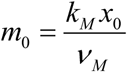

and dimensionless ratios

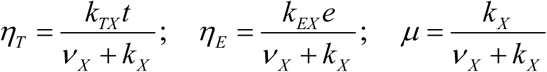

with which they become

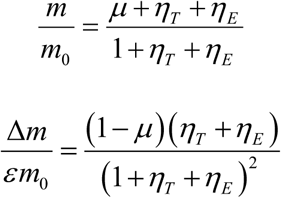

where again we take *ò_T_* =*ò_E_* ≡ *ò* as above. These expressions are reproduced in Eq. 2 of the main text.

#### Parameter selection in the mathematical models

In the mutual repression model, we sample the ratios *η_X_* , *η_Y_* , *μ_T_* , *μ_E_* , *β_T_* , and *β_E_* logarithmically in the range [10^−3^ 10^3^ ] and accept each sample if Δ*m* is smaller than (i) its value with only the TGF-β gradient ( *μ_E_* = *β_E_* = 0 ) and (ii) its value with only the EGF gradient (*μ_T_* = *β_T_* = 0 ), (iii) and both i and ii are positive. The result, averaged over 10^7^ samples, is shown in Figure. 4C and D.

In the shared pathway model, we sample the ratios *η_T_* and *η_E_* logarithmically in the range [10^−3^ 10^3^ ] and *μ* uniformly in the range [0 1], and accept each sample if Δ*m* is smaller than (i) its value with only the TGF-β gradient (*η_E_* = 0 ) and (ii) its value with only the EGF gradient (*η_T_* = 0 ), (iii) and both i and ii are positive. The result, averaged over 10^7^ samples, is shown in Figures. 4F and G and 5A and B. ɛε

### Derivation of the phase diagram in Figure. 6A

The boundaries in the phase diagram in Figure. 6A follow from the expression for Δ*m* in Eq. 2 of the main text as

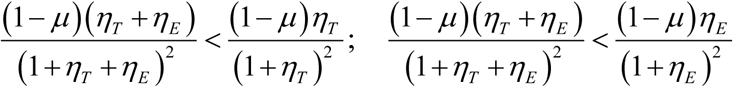

where the righthand sides are the cases with only the TGF-β gradient (*η_E_* = 0 ) and only the EGF gradient (*η_T_* = 0 ), respectively. If both conditions are true, it is antagonism; if neither is true, it is synergy. Defining the amplification factor and relative pathway strength

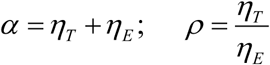

as in the main text, or equivalently

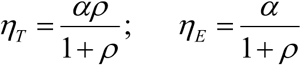

the conditions become

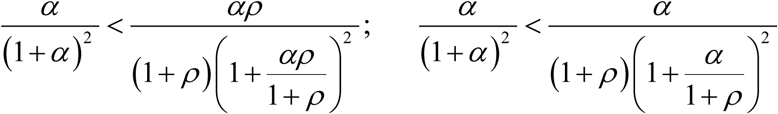

which simplify to

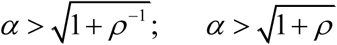

These curves are plotted in Figure. 6A.

## QUANTIFICATION AND STATISTICAL ANALYSIS

In evaluating the chemotactic characteristics, more than 30 trajectories were analyzed in an experimental trial, which was repeated at least three times for all experimental groups. Each trajectory produced a value of CI and speed. All collected CIs and speeds in each group were reported in the box plots. A data point in the box plots indicates each metric of a cell trajectory showing the distribution characteristics of the metric in a group. Differences in median CIs and speeds presented in the box plots were statistically analyzed by Mann-Whitney U-test. The significant changes between comparisons were examined when the p value <0.05. To evaluate the chemotactic accuracy and cell motility, medians of CIs and speeds from the repetitions (n>3) were averaged and reported in a bar with an error bar representing a standard estimated error (S.E.). Differences in CIs presented in the bar graphs were statistically analyzed by student t-test. The significant changes between comparisons were examined when the p value < 0.05.

**Movie S.1 Time-lapse of cell migration, Related to Figure1.**

## SUPPLEMENTAL INFORMATION

**Figure. S1.**
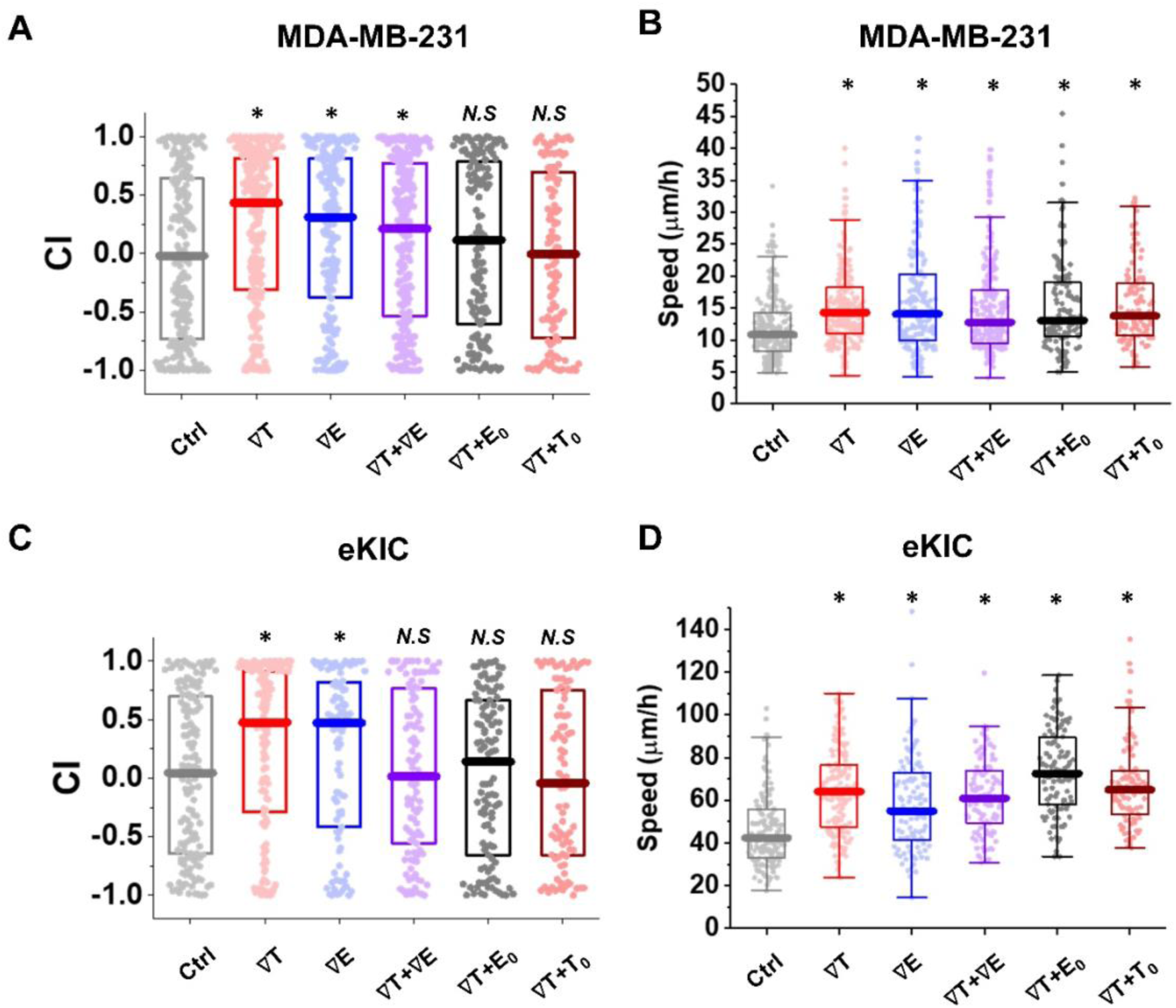
CI and Speed of all collected trajectories, Related to Figure2, Figure3, and Figure 5. (A) Distribution of chemotactic index (CI) from all trajectories of MDA-MB-231 collected from experimental trials (N>3) including >100 trajectories per condition, respectively. Box represents quartiles with a median line in the middle of the box. (B) Distribution of speed from all collected trajectories of MDA-MB-231. Box: interquartile range (IQR) ± 1.5 IQR whiskers with a median line. (C) Distribution of chemotactic index (CI) from all trajectories of eKIC. (D) Distribution of speed from all collected trajectories of eKIC. A dot represents data from a single trajectory. (*: p<0.05 for comparison with ctrl, N.S: no significance with p>0.05 in Mann-Whitney test)

**Figure. S2.**
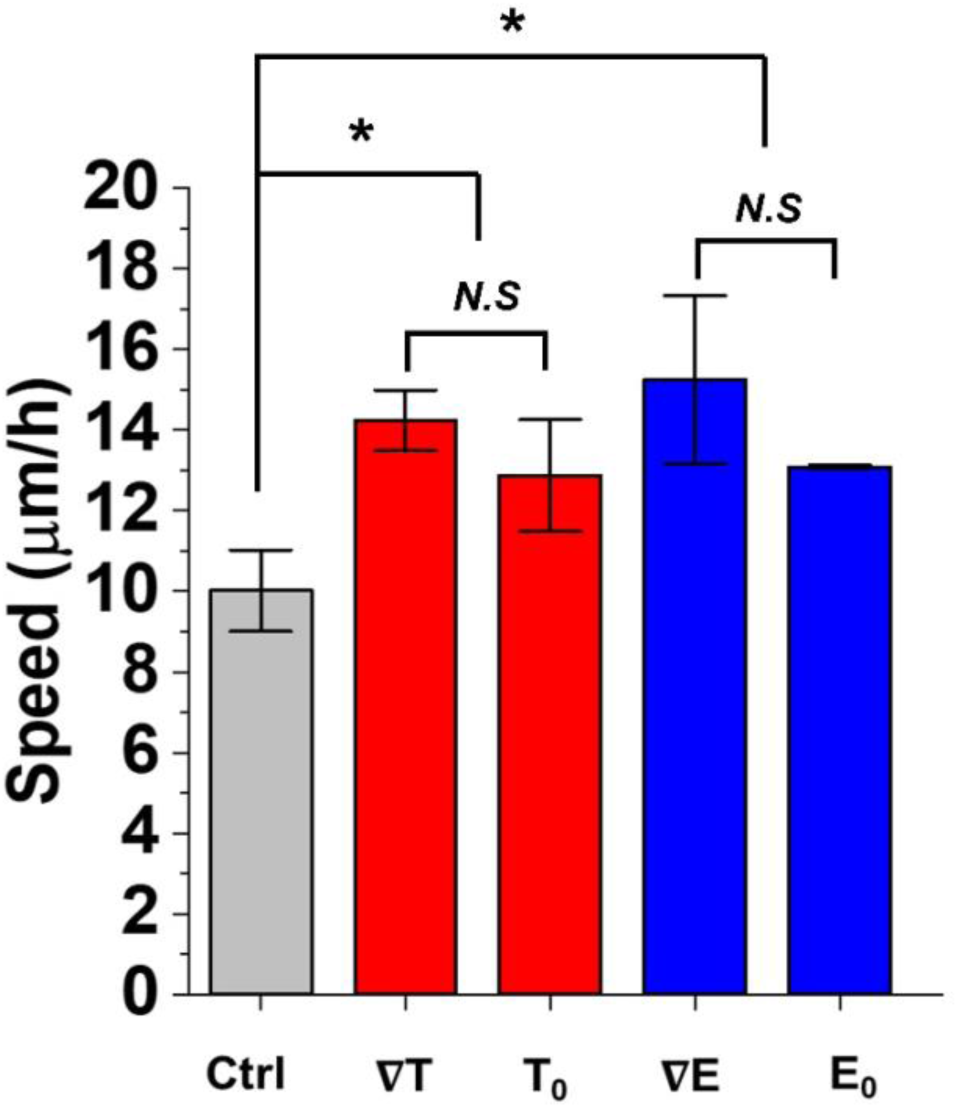
Speed of MDA-MB-231 in the uniform background concentration of corresponding gradient, Related to Figure 4. Control, 50nM/mm TGF-β gradient (∇T), 25nM uniform TGF-β (T_0_), 800nM/mm EGF gradient (∇E), and 400nM uniform EGF (E_0_). Speed of all treatment groups (∇T, T_0_, ∇E, and E_0_) increase compared to Ctrl, while T_0_ or E_0_ is not significantly different from the corresponding gradient (∇T or ∇E). Bar represents mean of the medians ± S.E (n≥3). The medians are collected from >35 trajectories in a sample, respectively. (*: p<0.05, and N.S: no significance with p>0.05 in student’s t-test)

**Figure. S3.**
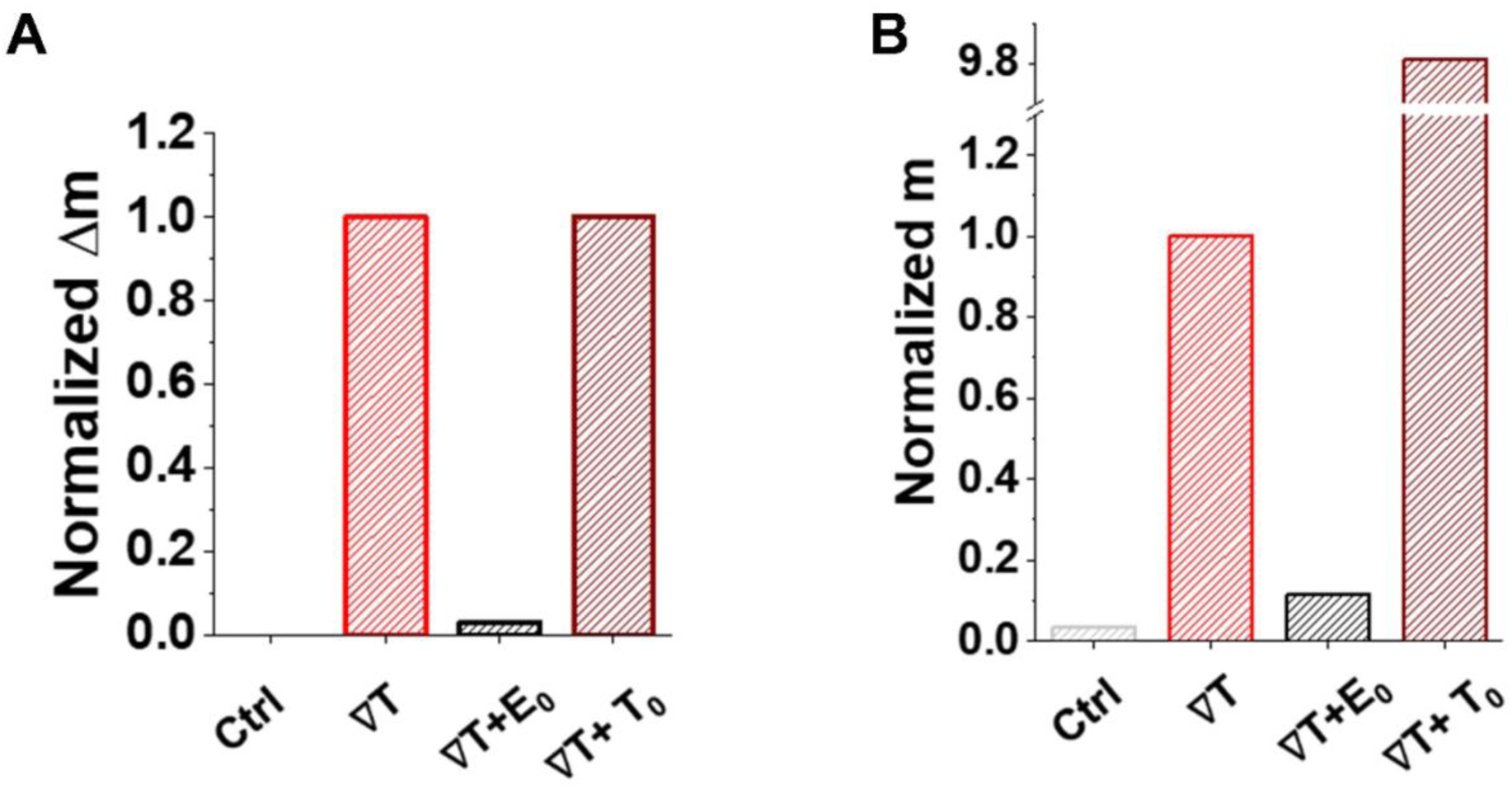
The crosstalk model prediction, Related to Figure 5. (A) The bias Δ*m* decreases when a TGF-β gradient ∇T is combined with a uniform background of EGF (∇T+E_0_), but not with a uniform background of TGF-β (∇T+T_0_). The prediction is not comparable with the experimental results (Figure. 5C). (B) The average *m* decreases in ∇T+E_0_, and significantly increases in ∇T+T_0_, which is a different tendency from the experimental results (Figure. 5D) showing comparable speeds each other when ∇T is combined with either ∇T+E_0_ or ∇T+T_0_.

